# Modeling the Opening SARS-CoV-2 Spike: an Investigation of its Dynamic Electro-Geometric Properties

**DOI:** 10.1101/2020.10.29.361261

**Authors:** Anna Kucherova, Selma Strango, Shahar Sukenik, Maxime Theillard

## Abstract

The recent COVID-19 pandemic has brought about a surge of crowd-sourced initiatives aimed at simulating the proteins of the SARS-CoV-2 virus. A bottleneck currently exists in translating these simulations into tangible predictions that can be leveraged for pharmacological studies. Here we report on extensive electrostatic calculations done on an exascale simulation of the opening of the SARS-CoV-2 spike protein, performed by the Folding@home initiative. We compute the electric potential as the solution of the non-linear Poisson-Boltzmann equation using a parallel sharp numerical solver. The inherent multiple length scales present in the geometry and solution are reproduced using highly adaptive Octree grids. We analyze our results focusing on the electro-geometric properties of the receptor-binding domain and its vicinity. This work paves the way for a new class of hybrid computational and data-enabled approaches, where molecular dynamics simulations are combined with continuum modeling to produce high-fidelity computational measurements serving as a basis for protein bio-mechanism investigations.

## 1. Introduction

Since its emergence in December 2019, the novel severe acute respiratory syndrome coronavirus 2 (SARS-CoV-2) continues to be a major concern due to its high transmission rate and worldwide presence [2, 41, 42, 44]. The mechanism by which the virus interacts with cells is mediated by the receptor-binding domain (RBD) on the SARS-CoV-2 spike protein (S protein). The full S protein, composed of three identical S proteins to form a homotrimer, binds to the human angiotensin-converting enzyme 2 (ACE2), and this interaction is the principal instrument for viral infection [2, 18, 20, 33].

Multiple studies maintain that the spike protein’s structure and fxunction are two features highly responsible for cell infection because the conformational rearrangement of the S protein reveals its binding interface [15, 18, 37, 39]. Recently, Zimmerman *et al*. [46] produced the first molecular dynamics simulation of the spike opening. This computational *tour de force* involved millions of citizen scientists collaborating through the Folding@home initiative [1], and produced an overall 0.1s of molecular trajectories. For a full characterization of the protein interaction, the molecular trajectory may not be enough.

One aspect that may provide some insight into the interactions of the S protein is the electrostatic potential it generates. Indeed, according to a previous study, the affinity constant for the RBD of SARS-CoV-2 to the ACE2 is 10 to 15 times greater than that of SARS-CoV, potentially contributing to its transmission efficiency [37]. The reason for the higher binding affinity was attributed to several mutations, most notably from the residue Val404, found in SARS-CoV, to the positively charged Lys417 in SARS-CoV-2. This mutation resulted in an intensified electrostatic potential complementarity between the negatively charged ACE2 binding site and the now more positively charged RBD of SARS-CoV-2 [15, 37]. To understand the contribution of charges, and also to help inform drug design strategies that leverage this distribution, it is imperative to understand how the S protein electrostatic potential changes during its opening.

Although the electrostatic surface between the closed and open conformations of the spike protein is well characterized, all of the aforementioned work examine static structural data [2, 15, 37]. There is a deficit of numerical quantification regarding the surface of the entire S protein, the electrostatic potential on and around the protein’s surface, and how these properties vary as the protein transitions from a closed to an open conformation.

In this work, we leverage the S protein opening trajectory created by the Folding@home initiative to examine and characterize the electrostatic potential of the SARS-CoV-2 S protein during trimer opening. The novelty of our exploration lies in the combination of molecular dynamics simulation with partial differential equation modeling to obtain dynamical electric potential maps, thus revealing variations of the electro-geometric properties of the S protein. At each frame of the simulated trajectory, we reconstruct the S protein surface and calculate the generated electric potential with the approach developed by Mirzadeh *et al*. [26, 27]. Using adaptive non-graded Octree grids and sharp discretizations we efficiently produce high-fidelity solutions to the non-linear Poisson-Boltzmann equation. In the continuum model, all temporal variation may be neglected, and as a consequence, all potential maps are independent, making their computation parallelizable.

Our study of the electrostatic dynamics of the S protein opening reveals dramatic rearrangements of the electrostatic field during this process. These rearrangements act to localize a negatively charged field towards the interior of the S protein and expose a positive surface of its residue binding domain. This is in line with the negative charge of the target binding region on the ACE2 receptor and may aid the S protein in binding to its target with high affinity.

The paper is arranged as follows: in section 2.1, we present our molecular trajectory emphasis and investigate the spike protein’s conformational changes. The dynamical electrostatic and geometric properties of the spike protein are characterized in section 2.2, followed by an analysis and fundamental takeaways in section 3. Section 4.1 entails a thorough mathematical design for the calculation of the potential field that was carried out on each frame in the trajectory. Section 4.2 encapsulates the numerical method used accompanied by a convergence study for computational validation.

## 2. Results

### 2.1. Opening of the SARS-CoV-2 Spike Protein

The S protein exists primarily in the closed configuration, hiding the three identical RBDs in its core [46]. The Folding@home trajectory focuses on the opening of the unglycosylated, uncleaved spike configuration through the detachment of a single monomer from the spike’s core, revealing its associated binding site. A similar (but not identical) monomeric-opening state has recently shown to be populated in ≈ 16% of the unbound spike population in a recent cryoEM study [5].

#### 2.1.1. Simulated Trajectory

The trajectory, from Zimmerman *et al*. [46] contains a 71-frame sequence that captures the most populated pathway of the spike protein’s transition from its closed to open state. This path was calculated by means of a goal-oriented adaptive sampling algorithm (FAST, [45]) to favorably sample spike opening.

Each frame consists of a list of *N* = 51, 671 atoms, represented by their positions **x**_*i*=1..*N*_, radius *r*_*i*=1..*N*_ and fixed partial charge *z*_*i*=1..*N*_. Throughout these frames, the protein undergoes continuous deformations which shift the receptor-binding domain of one of the monomers from being hidden, while the spike is in its closed conformation, to being fully exposed once the spike opens. Figure 1*a* and 1*b* represent the protein’s molecular structure in the open and closed configurations. Each one of its chains is depicted in a different color to illustrate the protein’s trimeric structure. The revealed receptor-binding domain is located at the opening extremity of the red chain (see Fig. 1*a*,1*b*).

**Figure 1:**
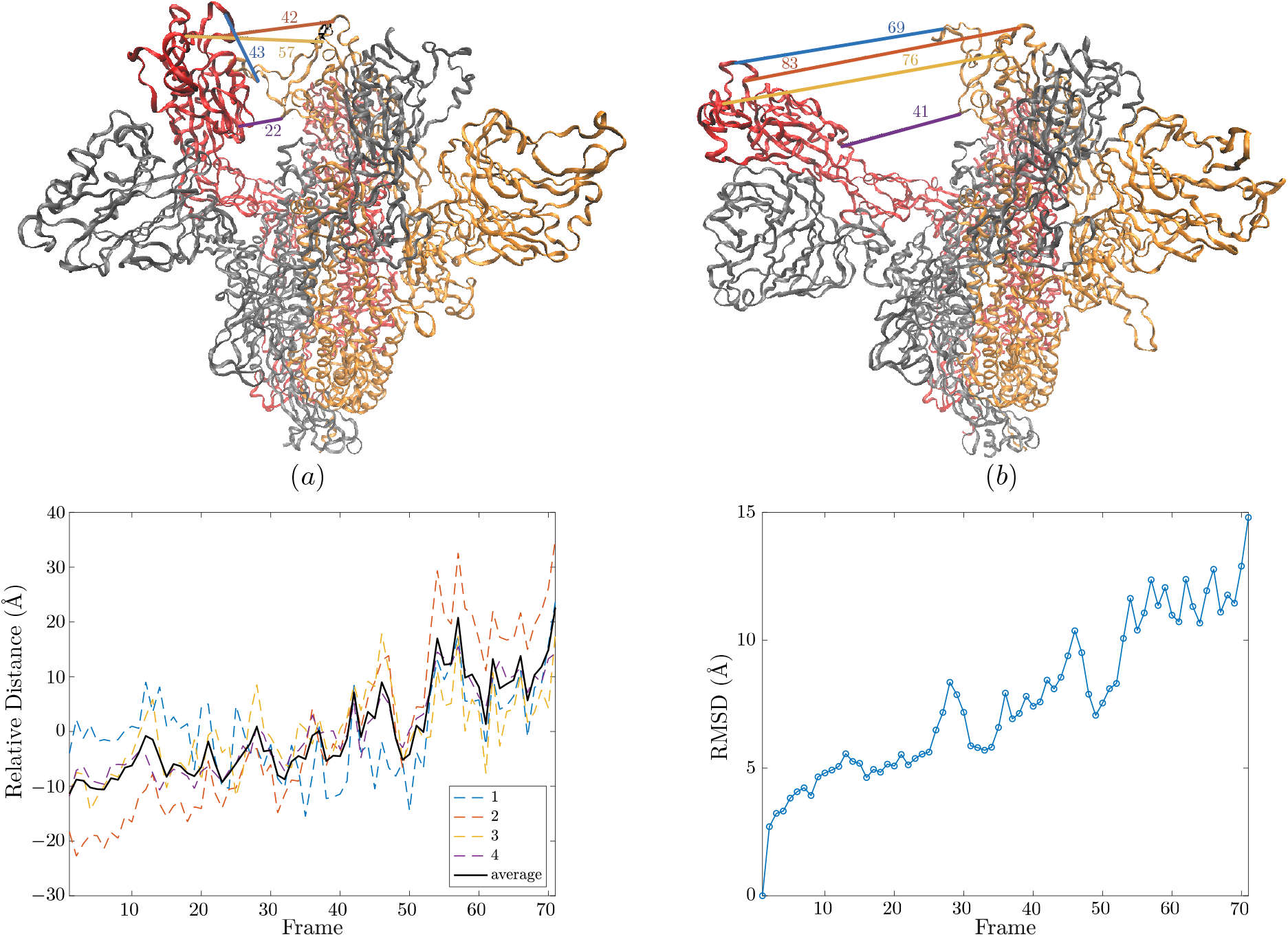
SARS-CoV-2 Spike protein visualized in its closed (*a*) and open (*b*) states. The four pairs of atoms, along with the separating distances, used to quantify the gap opening are represented with straight solid colored lines. (*c*) The relative distance (*Å*) between arbitrarily chosen atom pairs is represented as a function of time. One atom is selected on the receptor-binding domain (RBD) and the other opposite the RBD across the opening of the spike protein. This distance is relative to the average distance between each respective atom pair (1,2,3, and 4 in the graph), hence the term ‘relative distance’. (*d*) The root mean squared deviation (RMSD) is calculated as a function of time with frame 1 (closed state) as the reference frame.

#### 2.1.2. Relative Opening Measurement

To provide preliminary quantification of the spike opening, dozens of atom pairs on opposing sides of the spike were chosen and the distance between them measured as the spike transitions from one conformation to another (i.e. as a function of frame) (Fig.1*a* and *b*). One of the atoms in the pair was chosen from the binding interface, the monomer depicting in red in Fig.1(*a* and *b*), while the second atom in the pair was chosen across the top of the spike opening. Although all pairs presented the same general trajectory, four of these atom pairs were arbitrarily chosen as a subset for illustration.

The resulting relative distance between the atom pairs is measured in Angstroms (Å), as shown in Fig. 1*c*. The lines labeled 1, 2, 3, and 4 refer to the behavior of the four-pair subset, with their average behavior depicted in black. All four measurements generally follow the same trend. This extension happens continuously outside of frames 50 to 55 which show abrupt variations.

#### 2.1.3. Magnitude of the Conformation Change

To measure the total structural deformation we compute the root-mean-square deviation (RMSD). It measures the average distance between atoms in the current position and some reference configuration, defined here as the initial closed configuration. Specifically

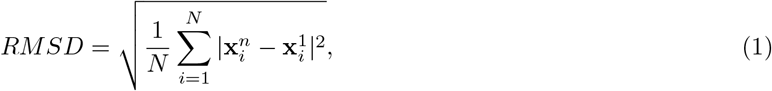

where the superscripts *n* and 1 are for the current and initial states. Despite large variations at the initial and final stages, the RMSD evolution (depicted in Fig. 1*d*), shares similar features with the evolution of the relative opening. These similarities suggest that the significant transformation that occurs during the opening are recapitulated by the 4 distances in Fig. 1*c*.

### 2.2. Computational Investigation

The S protein remains predominantly in the closed conformation to mask its receptor-binding domains (RBDs), thereby impeding their binding. To bind with ACE2, the S protein transforms into its open conformation, revealing its binding interface [46].

In describing our results, we refer to the part of the spike containing the three RBDs as the top part, and the predominantly negatively charged portion binding to the virus capsid as the bottom part.

Our simulations (see Fig. 2*a*) illustrate that the top part of the spike protein is predominantly positively charged, in accordance with the negative charge of the ACE2 binding site (see Fig. 3*a*). Surprisingly, the top of the spike also reveals an underlying core with a surface area that has a dense negative charge. As the spike opens, this area could be exposed to the solvent, generating a negatively charged electrostatic cloud in the upper part of the protein, which may repulse the negatively charged ACE2 receptor. Therefore, we wanted to characterize the dynamics of the geometrical and electric properties of this negatively charged core *χ* (Fig. 2*b*), the resulting repulsive cloud 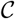 (Fig. 2*c*), the binding site 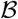 (Fig. 3*c*), and the far electric field of the spike protein as they shift from the dynamics that occur during spike opening.

**Figure 2:**
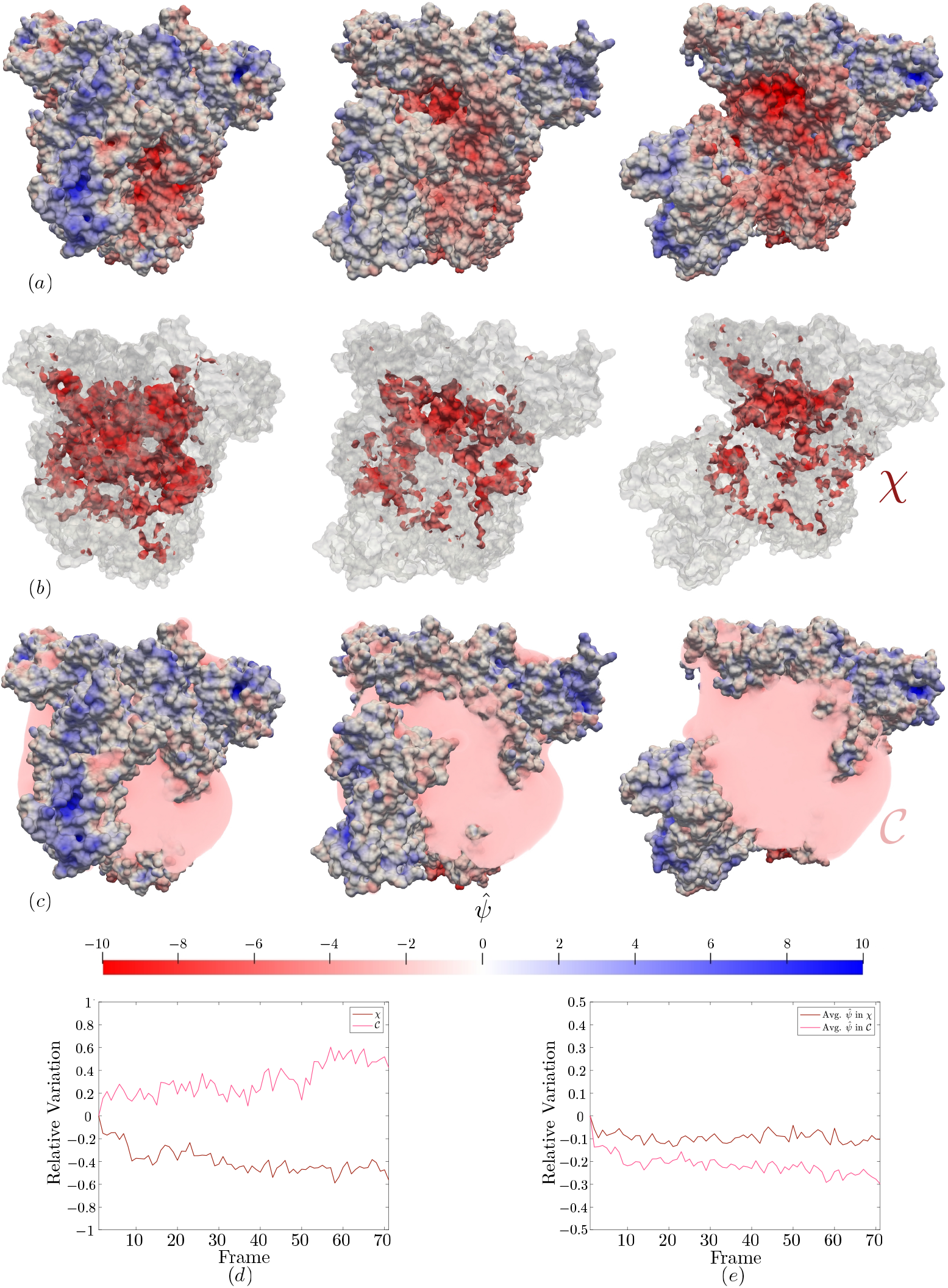
(*a*) Potential map of the SARS-Cov-2 spike open (frame 1), intermediate (frame 35), and close configuration (frame 71). (*b*) Evolution of the negatively charged core as the spike opens. (*c*) Repulsive electric cloud generated by the negatively charged core. (*d* − *e*) Characterization of the negatively charged core *χ* and repulsive electric cloud 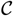: relative size (*d*) and potential (*e*) variations over the entire molecular trajectory.

**Figure 3:**
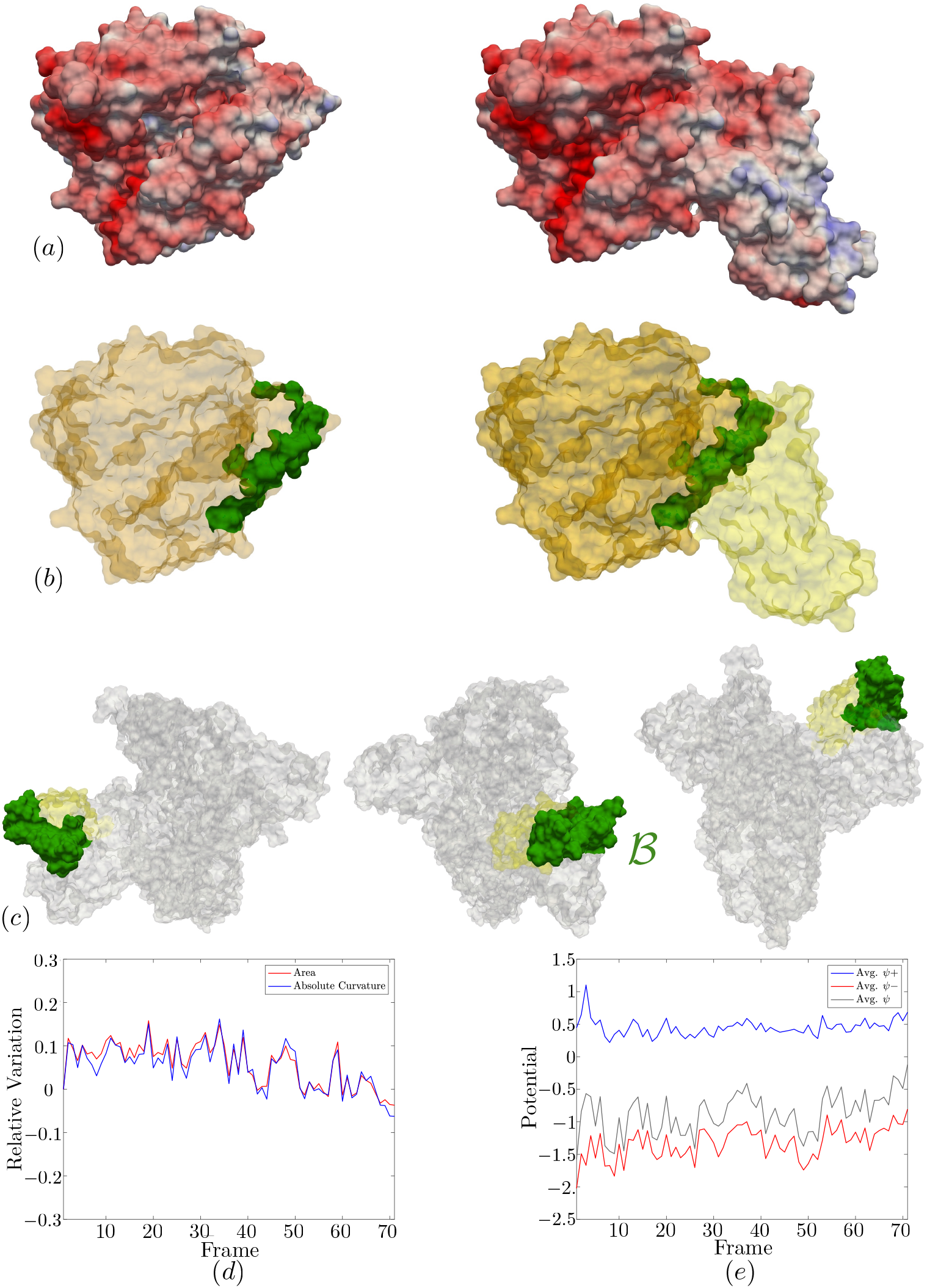
(*a*) Portion of the ACE2 receptor displaying surface charge (left). ACE2 and RBD of the spike protein connected (right). (*b*) Binding site of ACE2 highlighted in green (left). The binding site of ACE2 is highlighted within the ACE2 (orange) and RBD (yellow) connection (right). (*c*) SARS-Cov-2 spike in open conformation with RBD (yellow) and binding site, B (green) highlighted. (*d* − *e*) Characteristics of the binding site, 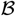, of the spike protein. (*d*) Relative variance of the area and absolute curvature along with average potential. (e) average positive and negative potential.

#### 2.2.1. Negatively Charged Core

We defined the negatively charged core *χ*, illustrated in Fig. 2*b*, as the area of the SES in the top of the protein where the electric potential is more negative than a threshold value *c_χ_* = −5. Figure 2 *c* and *d* depict the evolution of its surface and average charge. As the spike opens, the area shrinks by about 50%, while the variation in the average potential remains small (≈ 10%). In both cases, the most significant variations happen during the first 10 iterations. In comparison, the molecular structure analysis (see Fig. 1) displays continuous variation over the entirety of the trajectory, indicating that the structural and electro-geometric transformations are non-trivially coupled. The diminution of both quantities could be explained by the fact that the spike opening removes a barrier, one of the monomers, between the core and solvent, causing the dilution of the surface charges in the solvent, and therefore a contraction of the core and its charge.

#### 2.2.2. Repulsive Electric Cloud

We define the negatively charged electric cloud 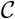, generated by the core *χ*, as the region in the upper part of the solvent where the potential is below the threshold value 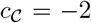 (see Fig. 2*c*). The measurements presented in Fig. 2*d* and *e* indicate 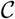 expands by ≈ 42% as the spike opens, while its average potential decreases in magnitude by 30%. Again we interpret this phenomenon as an effective dissolution of the negative charges induced by the removal of the protecting monomer.

For both geometries *χ* and 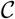, we observe large variations in the first 10 frames. The abrupt change (around frame 50) in the protein structure, discussed in section 2.1, is only perceptible in the size variation of the electric cloud 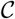.

#### 2.2.3. Binding Site Characterization

The interface between the SARS-CoV-2 receptor-binding domain (RBD) and ACE2 are of particular interest as the binding of the two facilitates virus entry into cells [2, 18, 20, 33]. Lan et al. [18] undertook the task of determining the residues of the RBD and ACE2 interface which form a connection, finding the location of the binding site on the spike, 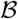, to consist of 17 residues between Lys417-Tyr505 and the ACE2 site to be formed of 20 residues between Gln24-Tyr83 and Asn330-Arg393.

For this analysis we define the binding site, 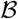, for the spike and the ACE2 receptor as the portion of the proteins SES generated by the residues sequences [417–505] and [24 – 42] U [79 – 83] U [330] U [354 – 357] U [393] respectively. The spike RBD is generated by the sequence [333-527]. When the spike is in the closed position 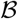 is located near its center, but as the spike shifts to its open conformation, 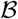 is moved outward and exposed to the solvent, as seen in Fig. 4. As the spike opens, we monitor the evolution of the binding site area, average absolute curvature, and potential. The average is computed over the binding site surface. We interpret the average absolute curvatures as measures of the global convexity of the binding site. For the potential evolution, we distinguish between the average potential, average positive potential, and average negative potential.

**Figure 4:**
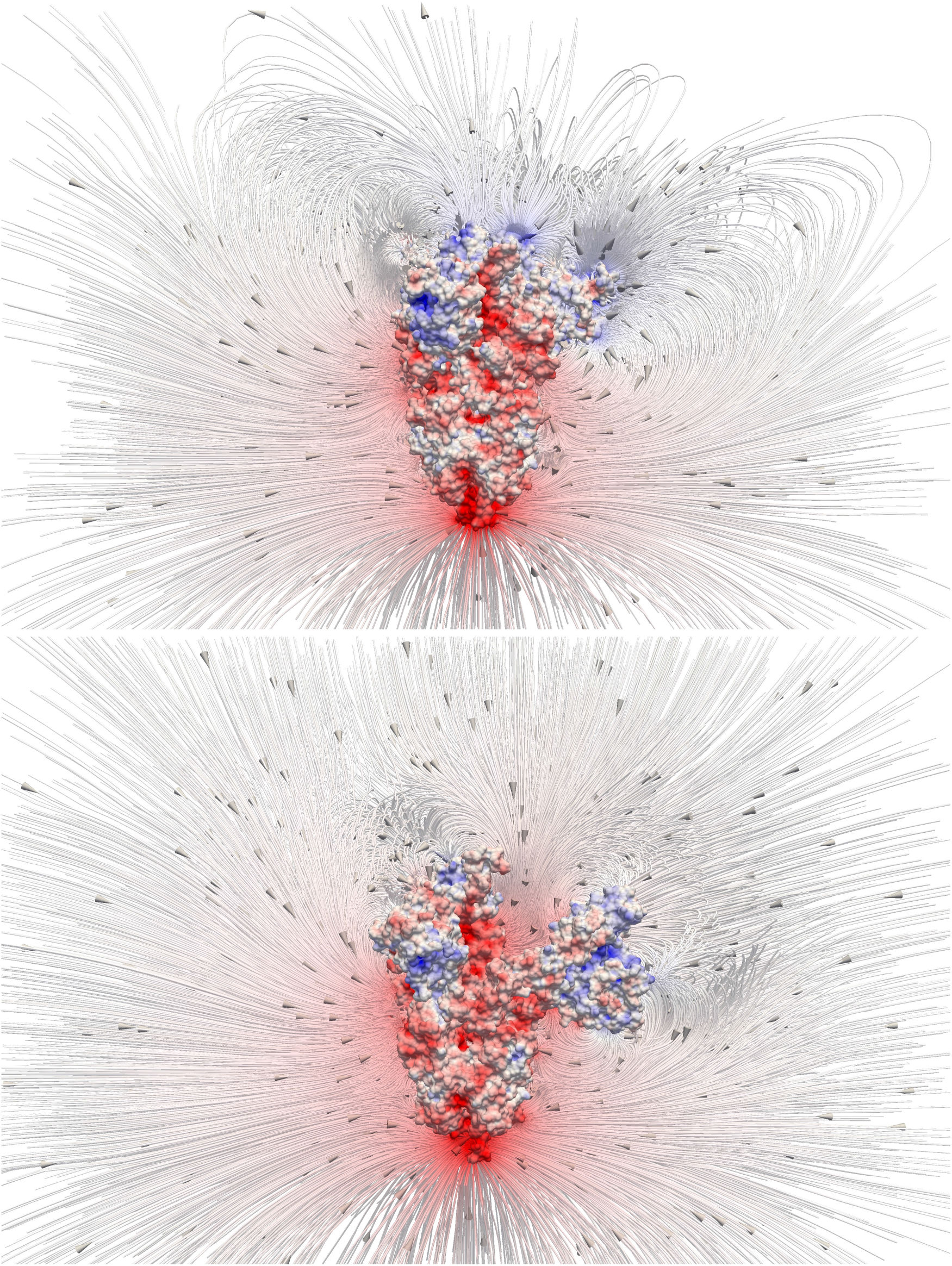
Streamlines and direction of the electric field in the close (top) and open (bottom) configurations. As the spike opens, the streamlines emanating from the upper part of the molecule are observed to open as well. This effect is particularly noticeable for the lines emerging from the RBD domain (opening extremity on the right).

Figure 3 provides a depiction of the resulting information. A relative decrease is observed for the area and average absolute curvature: as the spike opens the binding site shrinks and flattens, in each case by 10 – 15%, suggesting minor conformational changes. The average negative potential also experiences a decrease, becoming less negative as the spike opens. Meanwhile, the average positive potential remains fairly constant, resulting in the total average potential of 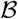 following very closely with the changes in average *ψ*^−^ and ultimately nearing 0 in the final frame. This increase in average potential is consistent with the knowledge that the ACE2 receptor it is designed to bind to is negatively charged, and may be important in directing the ACE2 binding to the correct region.

#### 2.2.4. Electric Field Structure

The spike electric far-field streamlines are depicted in Fig. 4. In the closed configuration, the majority of streamlines emanating from the bottom part of the spike are pointing away from the spike. Only a few of them, caused by the rare presence of positive charges, return rapidly to the protein. In the upper part, the streamlines are predominantly closed, suggesting that in this configuration the spike protein may not be able to attract charges of any polarity at long range, and therefore has limited attracting potential. This may cause a lower affinity. As the spike opens, the streamlines in the bottom remain unchanged. However, at the top of the protein, the streamlines are now predominantly open.

Fig 5(a)-(b) depicts the entire electric field structure along with the region in the solvent where the electric field point away from the protein, or equivalently where negative charges would be drawn to the spike. In the closed configuration, this region is predominantly located above the spike, while in the open configuration it is split into three sub-regions, each centered around one of the RBDs. In the latter, the sub-region centered around the revealed RBD represents 45% of the total region and carries 34% of its total electric potential energy.

**Figure 5:**
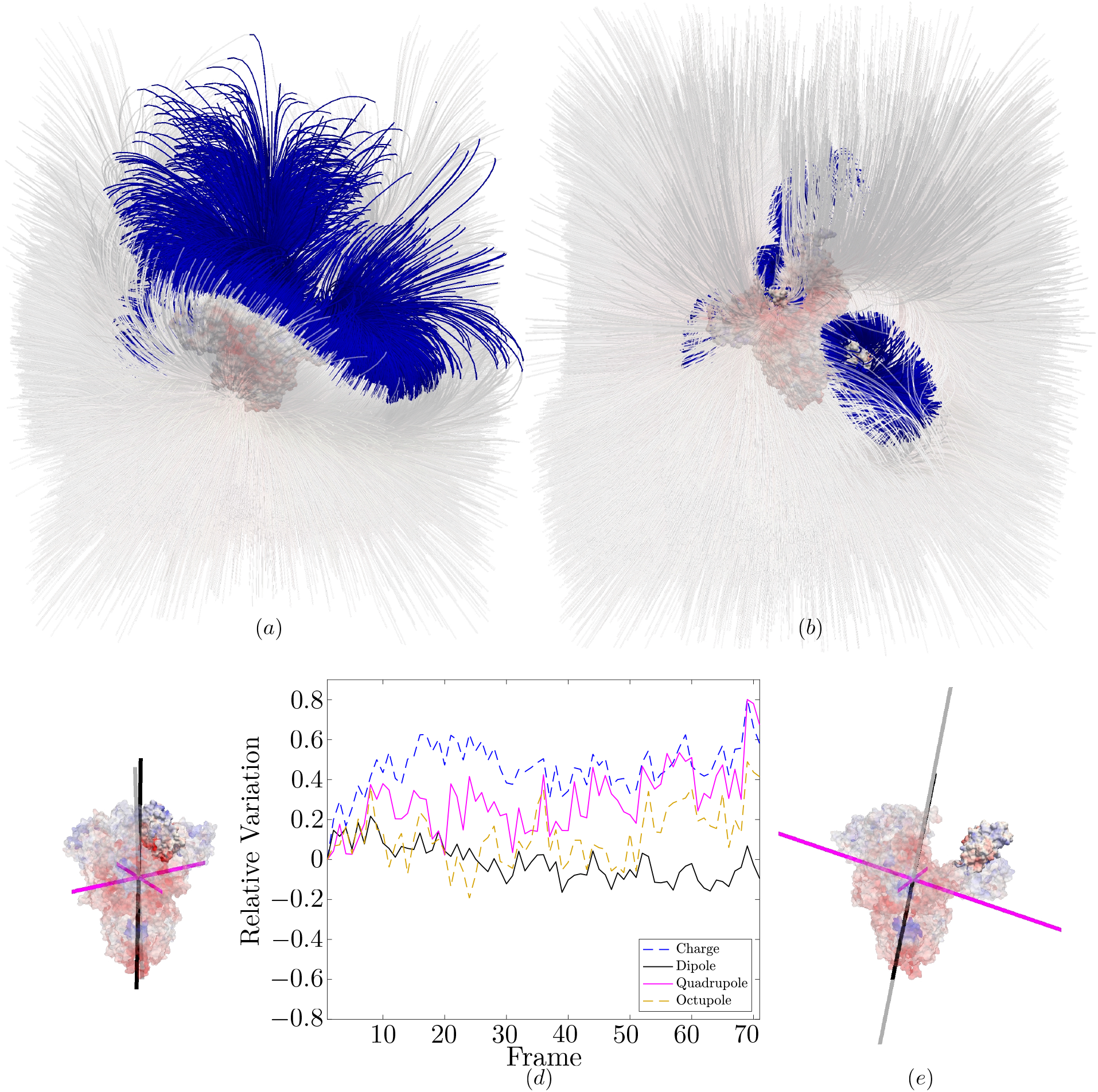
Streamlines of the electric field 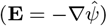 in the close (*a*) and open (*b*) configurations over the entire computational domain. The portion of the streamlines where the electric field is pointing away from the protein (i.e. ∇*ψ* · **x** = −**E · x** < 0), and therefore negative charges would be drawn to it, is depicted in blue. (*c* – *e*) Electric potential multipole structure: charge density decomposition over its first four moments *C, D, Q* and *O*. (*c*) and (*e*) represent the principal directions of the dipole *D* (in black) and the quadrupole *Q*, in the initial and final configurations respectively. For the quadrupole, the purple directions (eigenvectors) are associated with negative eigenvalues. The grey one has a positive eigenvalue. All segments lengths are proportional to the strength of the corresponding pole (*i.e*. the norm of the dipole or of the corresponding eigenvalue). The protein binding site is depicted as opaque. (*d*) depicts the evolution of the norm of all four first moments. We use the standard *L*^2^ entry-wise norm.

To characterize the restructuring of the electric field of the S protein, we define the first four moments of the non-dimensional charge distribution *q*(**x**) = sinh(−*ψ*(**x**)) in the solvent

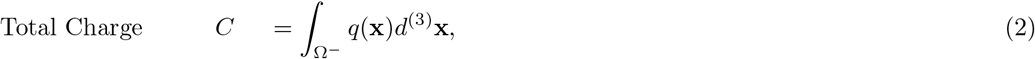

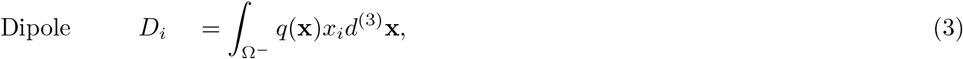

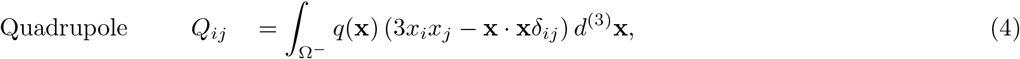

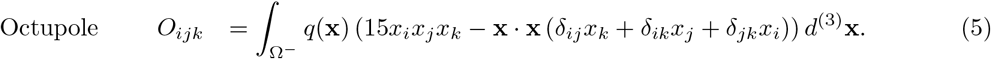

Note that the last two moments are defined in their symmetric traceless form, as they would be for a standard electrostatic multipole expansion. Because here our continuum model is more complex, it should be reminded that these four moments are not guaranteed to be the ones appearing in the multipole expansion. Nonetheless, they remain a pertinent tool to characterize the structure of the electrostatic solution. Their evolution throughout the opening, illustrated in Fig. 5c, show that the strength of the dipole (*D*) is decreasing, while all other moments norms are increasing. The two most informative moments for the structure of the electric field, the dipole and the quadrupole, are diminished by 9% and increased by 67% respectively. The principal direction of *Q* associated to the positive eigenvector appears to remain quasi-parallel to the dipole direction. In fact, all four directions (the dipole direction and the three principal directions of *Q*) appear to undergo the same rotation. The opening also reinforces the anisotropy in the tensor *Q*, by amplifying the difference between the magnitude of its eigenvalues, compressing one of its directions and effectively turning it into a two-dimensional quadrupole.

## 3. Discussion

The overarching objective was to reveal the impact of the SARS-CoV-2 spike protein’s transition from its open to closed conformation. Thus, we examined the entire protein’s electrostatic potential dynamics, focusing particularly on its receptor-binding domains. In the Folding@home trajectory we analyzed, one of the three monomers detaches from the core of the complex and becomes visible to the surrounding environment, consistent with recent cryo-EM derived structures [5]. Our results show that despite the dramatic molecular displacement engendered by this strategic re-positioning, the geometric and electric properties of the RBD itself remain largely unaltered. Rather, as we describe below, changes emanating from the exposure of the core of the S protein cause a change in the electric fields surrounding the RBD.

Our continuum analysis, both of the surface potential and the volumetric potential in the vicinity of the binding region point to the existence of an inner negatively charged core on the surface of the spike, which is revealed to the solvent as the spike opens. This negatively charged surface shrinks as the spike opens, inducing a negatively charged region between the exposed RBD and the two hidden ones. The emergence of this electric cloud is *a priori* puzzling: the binding site of the ACE2 receptor being almost entirely negatively charged, we would expect this cloud to repulse the receptor. However, this cloud does not envelop the spike’s binding site, 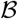, which becomes more positively charged. This is in line with what we expect is a strong binder of the ACE2 receptor, and indicates that targeting the open state of the S protein may be a more viable drug design strategy than the closed configuration. Indeed, recent cryoEM studies indicated that ≈ 16% of a recombinantly expressed S protein population is in an open state, with a single monomer “erected” from the core [5].

The electric field, which we observe uncoiling above the protein, exhibits a dramatic rearrangement. This is correlated to the emergence of the negatively charged cloud and manifests in the multipole moments decomposition. In particular, we observe the quadrupole moment growing in magnitude, rotating, and compressing one of its principal directions. This transformation can be interpreted as a strategic transition from an undirected configuration where the entire top part of the spike is attracting negatively charged structures, such as the ACE2 binding site, to one where the attraction is directed toward a specific RBD.

At the computational level, we have developed a novel framework, combining molecular dynamics simulations with a continuum physics numerical solver to produce high-resolution multiscale dynamical potential maps of deforming proteins and their surrounding environment. Our framework includes practical mathematical tools to quantify the numerical error and characterize protein electro-geometric properties. The molecular trajectories alone are insufficient for understanding protein electrostatic interactions and thus, protein core bio-mechanisms. On the other hand, it is unrealistic to envision an approach where trajectories of such a large protein could be obtained solely through continuum modeling. We believe hybrid approaches, such as the one presented here, can provide the scientific community with invaluable information, which we hope can be used to elucidate how changes to physical properties such as electrostatics translate into function.

The progression of time will uncover a multitude of breakthroughs regarding the behavior of the SARS-CoV-2 S protein. In the short time since the conception of our investigation, numerous discoveries by other researchers have been made public already. For example, the trajectory presented in [46] contains glycans, chemical compounds known to coat the exterior of many viruses, which may have an impact on the results presented here. Recall that the trajectory we explored here is simply one possible path the spike may follow, and so there is a possibility that a different trajectory would be a more accurate representation of the true function of the spike protein. Recent studies displayed cryo_EM structures for the spike, presenting an open configuration that differs from the open configuration represented here [40, 36]. A similar analysis to the one presented here may be required on different structures, as more probable trajectories are predicted and novel molecular structures uncovered, to capture a more relevant image of the electric potential on and around the spike. Utilizing the same scientific pipeline, we are confident high-fidelity quantitative insights — into the electric potential of molecules unrestricted to the one investigated here — could be obtained efficiently, supporting the global scientific effort.

## 4. Method

### 4.1. Continuum Modelling

The Poisson-Boltzmann equation has long been recognized as the representation of choice to model electric potential. It has drawn significant interest from the computational community since the pioneering calculations of Warwicker and Watson in the early 1980s [38], which has lead to the production of a broad variety of numerical solvers [4, 3, 6, 7, 13, 21, 17, 22] and open source software [12, 16, 32], built over the traditional spectrum of numerical methods. Such tools are, for example, employed in the context of drug development and discovery, to calculate solvation free energies [10, 14, 28]. Using massively parallel architectures [11], these calculations can be carried out on considerably large proteins, such as the entire HIV-1 capsid (Protein database entry: 3J3Q, 4, 884, 312 atoms).

This section describes the reconstruction of the potential map at each iteration from the current protein structure using the non-linear Poisson-Boltzmann (PB) equation. Because of the way the trajectory has been obtained, all frames are independent. Thus, the potential maps are decoupled and can be computed separately. A more comprehensive model, such as the Poisson-Boltzmann-Nernst-Planck model[43, 25], would consider the diffusion of the ions and introduce time derivatives in the partial differential equations. Because at the atomic scale (L = 1Å), for typical diffusivities (*D* ≈ 10^−9^*m*^2^*s*^−1^), the ionic diffusion time scale 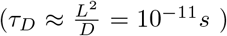 is orders of magnitude smaller than the estimated opening duration (*τ_O_* >> 10^−9^*s*), these effects can be neglected and the static PB equation is a pertinent model. Readers not interested in the numerical specifics can skip the following sections (4.1 and 4.2).

#### 4.1.1. Molecular Surfaces

A common way of portraying a molecular surface is by use of the van der Waals Surface depiction (see Fig. 6b). Each atom in a molecule is depicted by a sphere with location **x**_*i*=1…*N*_ and radius r_i=1_…_N_ that is defined by an isosurface on their electron density. The union of these spheres, depicted in gray in Fig. 6, forms the van der Waals Surfaces (vdWS) of that molecule. The nature of the vdWS means that some regions might be identified as being exposed to the solvent, while in fact, their geometry makes them inaccessible to the solvent particles. As a consequence, we define the Solvent Accessible Surface (SAS [19]) of the protein, depicted as the blue outline in Fig. 6 b; it is formed through the addition of the solvent’s particle radius, *r_P_*, to each *r*_*i*=1..*N*_ resulting in a buffer around the vdWS. While the puffed-up SAS depiction may be useful for some areas of study the Solvent Excluded Surface (SES) is preferable when discussing surface details of a molecule [31], as shown in Fig. 6b. As the name suggests, the inner molecule region defined by this surface includes all locations the solvent cannot occupy including the vdWS and the tiny crevasses on its exterior, shown as concave black triangles at the meeting of atoms A and B in Fig. 6b.

**Figure 6:**
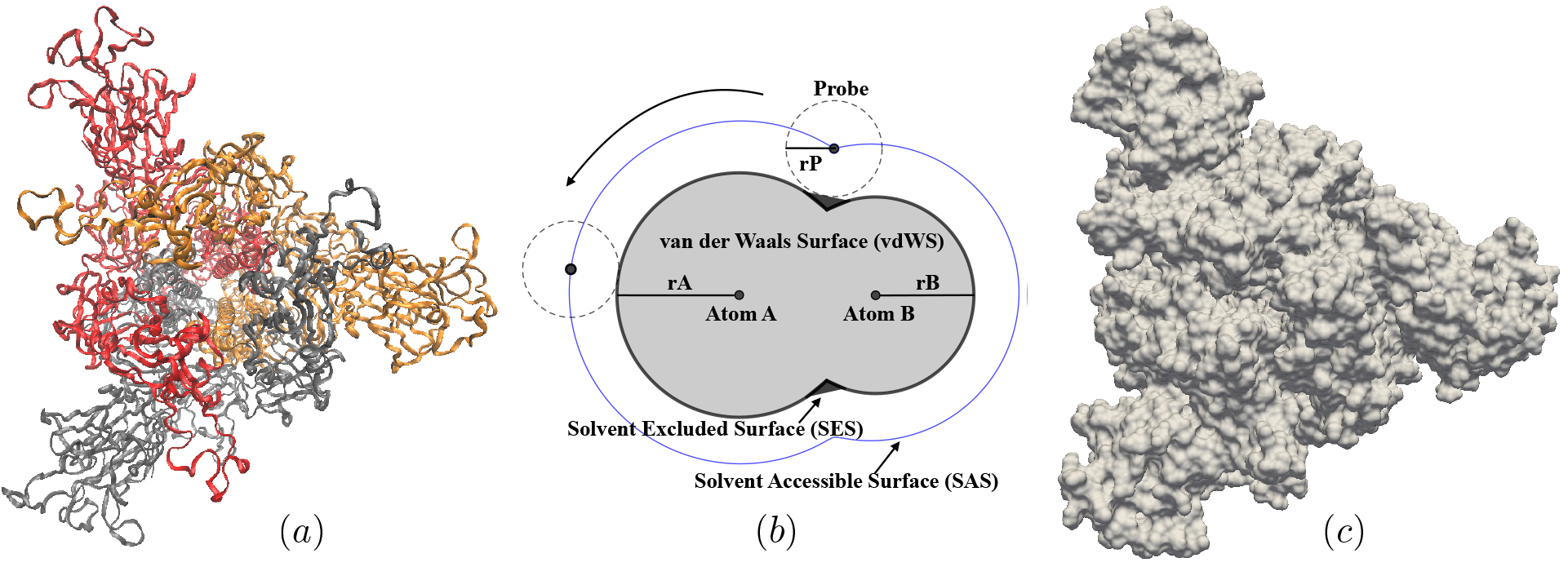
(*a*) Visualization of the top view of the Spike protein molecule (colored by monomer). Color denotes the three monomers (*b*) A cutaway diagram of a general molecular surface depicting the Van der Waals Surface (vdWS), solvent-excluded surface (SES), and solvent-accessible surface (SAS). (*c*) SES representation of the Spike protein oriented the same way as Fig. (*a*) (top view).

#### 4.1.2. Mathematical Representation and Numerical Construction

To represent the biomolecule, we employ the level set method [30] and capture the SES location as the zero level set of an auxiliary field *φ*(**x**) defined over the domain of interest Ω. The solvent Ω^−^, the Solvent Excluded Surface Γ, and the inside of the molecule Ω^+^ are defined as

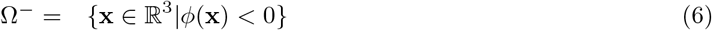

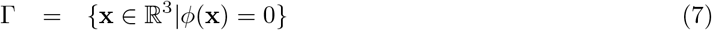

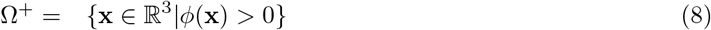

The normal **n** to the interface Γ is defined as pointing toward Ω^+^. It is calculated as the normalized level set gradient

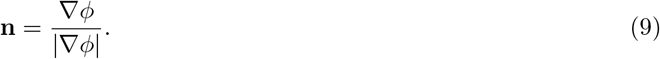

The level set function is constructed following the approach proposed by Mirzadeh *et al*. in [27]. We start by constructing the level set function *Φ_SAS_*(**x**) representing the SAS as

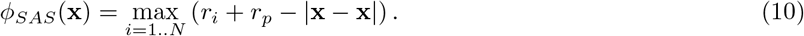

The SAS level set function is then reinitialized to be a signed-distance function (i.e. ∇ϕ_SAS_| = 1) by solving the reinitialization equation in fictitious time *τ* until the steady state is reached

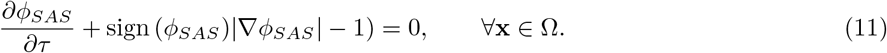

From the reinitialized function 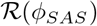, the SES level set function is then obtained as

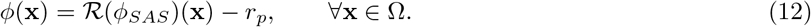

As it was pointed out in [27], this procedure can create non-physical inner cavities. They are identified by finding physical points disconnected from the contour of the computational domain *∂*Ω where the level set function *φ*(**x**) is positive. In practice, such points are isolated by solving the following Laplace problem

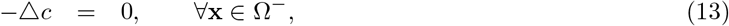

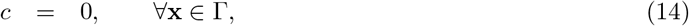

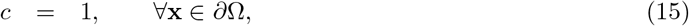

and detecting where *φ*(**x**) < 0 and *c* = 0. The cavities are removed by switching the sign of the level set function at these problematic positions. Finally, for computational purposes the SES level set is systematically reinitialized.

#### 4.1.3. Poisson-Boltzmann Equation

The electrostatic potential, Ψ, around a biomolecule immersed in a binary z:z electrolyte solution can be described by the following nonlinear Poisson-Boltzmann (PB) equation,

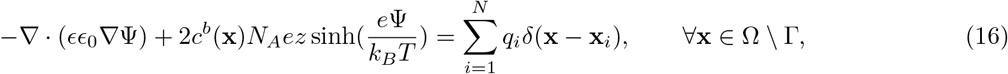

where *ϵ* is the relative permittivity, being equal to *ϵ*^+^ in the molecule (Ω^+^) and *ϵ*^−^ in the electrolyte (Ω^−^). *ϵ*_0_ is the permittivity of a vacuum, *N_A_* is the Avogadro number, *c^b^* is the bulk salt concentration, *e* is the elementary charge, *k_B_* is the Boltzmann constant, *T* is the temperature, *z* is the valence of the background electrolyte, *q_i_* and *x_i_* are the partial charge and position of the *i*^th^ atom respectively, *N* is the total number of atoms in the molecule. *c^b^*(*x*) = 0 inside the molecule. In non-dimensional form, the Poisson-Boltzmann equation takes the following form

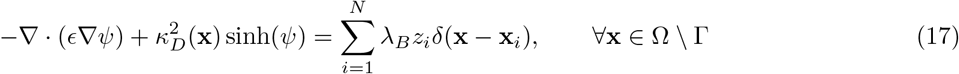

where the characteristic length *L* is chosen to be 1Å, the potential is scaled by the thermal voltage 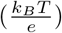, *z_i_* is the non-dimensional partial charge on the *i^th^* atom, and 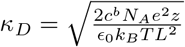 is the non-dimensional inverse of the Debye length. Inside the molecule, *κ_D_* is null. The constant 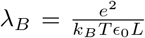 is the non-dimensional Bjerrum length. The above non-dimensional PDE is completed with the following jump conditions on non-dimensional potential

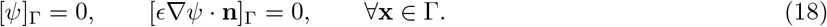

where the jump operator is defined for any quantity *ζ* defined in both domain as [ζ]_Γ_ = *ζ*^+^ – *ζ*^−^. All parameters and non-dimensional numbers, along with their values for this study, are summarized in table 1.

**Table 1:** Problem parameters and non-dimensional numbers.

| Parameter | Symbol | Definition | Value |
| --- | --- | --- | --- |
| Characteristic length | $L$ | - | $1\text{\AA}$ |
| Characteristic potential (thermal voltage) | $\psi_0$ | $\frac{k_B T}{e}$ | $2.570 \times 10^{-2} V$ |
| Vacuum permittivity | $\epsilon_0$ | - | $8.854 \times 10^{-12} Fm^{-2}$ |
| Spike relative permittivity | $\epsilon^+$ | - | 2 |
| Solvent relative permittivity | $\epsilon^-$ | - | $7.830 \times 10$ |
| Boltzmann constant | $k_B$ | - | $1.381 \times 10^{-23} JK^{-1}$ |
| Avogadro number | $N_A$ | - | $6.022 \times 10^{23} M^{-1}$ |
| Elementary charge | $e$ | - | $1.602 \times 10^{-19} C$ |
| Temperature | $T$ | - | $298.15 K$ |
| Salt concentration | $c^b$ | - | $1 \times 10.0^{-3} Mm^{-3}$ |
| Probe radius | $r_p$ | - | $1.4\text{\AA}$ |
| Valence of the background electrolyte | $z$ | - | 1 |
| Non-dimensional inverse of Debye length | $\kappa_D$ | $\sqrt{\frac{2c^b N_A e^2 z}{\epsilon_0 k_B T L^2}}$ | $9.207 \times 10^{-3}$ |
| Non-dimensional Bjerrum length | $\lambda_B$ | $\frac{e^2}{k_B T \epsilon_0 L}$ | $7.039 \times 10^3$ |

#### 4.1.4. Solution Decomposition

Following [27], we treat the singularities arising in the solution due to the singular charges inside the molecules by using the decomposition proposed by Chern *et al*. [8]. Doing so we split the non-dimensional potential *ψ*, into regular and singular parts: 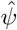 and 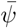, respectively

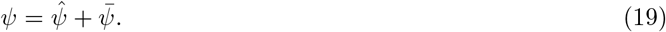

The singular part is itself split into two parts *ψ\** and *ψ*_0_

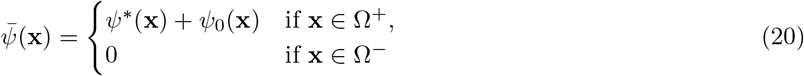

where *ψ*^*^ is the Coulombic potential due to singular charges,

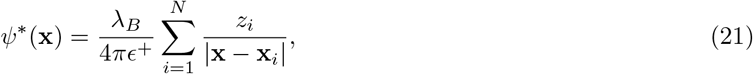

and *ψ*_0_ satisfies the following Poisson’s problem

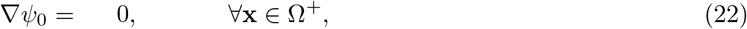

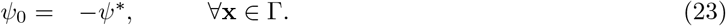

Utilizing the decomposition shown above, the regular (non-singular) part of the solution is given by solving

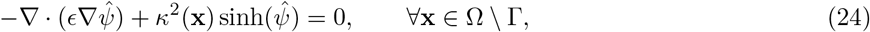

subject to the following jump conditions:

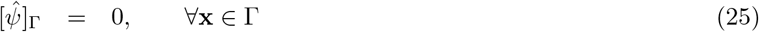

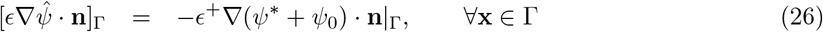

Since *ψ*\* is known analytically, the gradient, ∇*ψ*\*, appearing in the right hand side of (26) can be computed exactly

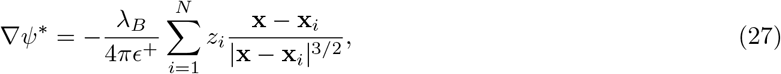

while ∇*ψ*_0_ must be numerically approximated.

### 4.2. Numerical Method

The numerical approach for the resolution of the above Poisson-Boltzmann problem is presented in this section, along with novel practical *a postieri* error estimates. We then verify the method for the entire trajectory by conducting a systematic convergence study for these estimators.

#### 4.2.1. Implementation

The numerical method, implemented on non-graded adaptive Octree grids, follows the general description given in [27], with each snapshot of the protein treated independently. The surface and grid generation is done using the level set framework developed by Min and Gibou [23]. In particular, as Fig. 7 illustrates, the mesh is systematically adapted to the SES location. All quantities are stored at the nodes of the mesh for improved accuracy and facilitated manipulation. The numerical solutions of the Poisson systems (13)-(14)-(15) and (22)-(23) (for the cavities detection and the construction of the regular part of the solution *ψ*_0_ respectively) are obtained using the second-order approach presented in [35], itself based on [24]. The solution, 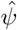, to the problem defined by Eqs. (24), (25) and (26) is constructed using a nodal version of the jump solver presented in [34]. The non-linearity of Eq. (24) is addressed using Newton’s method [8, 26, 27], with a relative error tolerance of 10^−6^, chosen to be orders of magnitude smaller than the desired overall numerical error. The whole method is parallelized in a shared memory fashion using OpenMP [9, 29].

**Figure 7:**
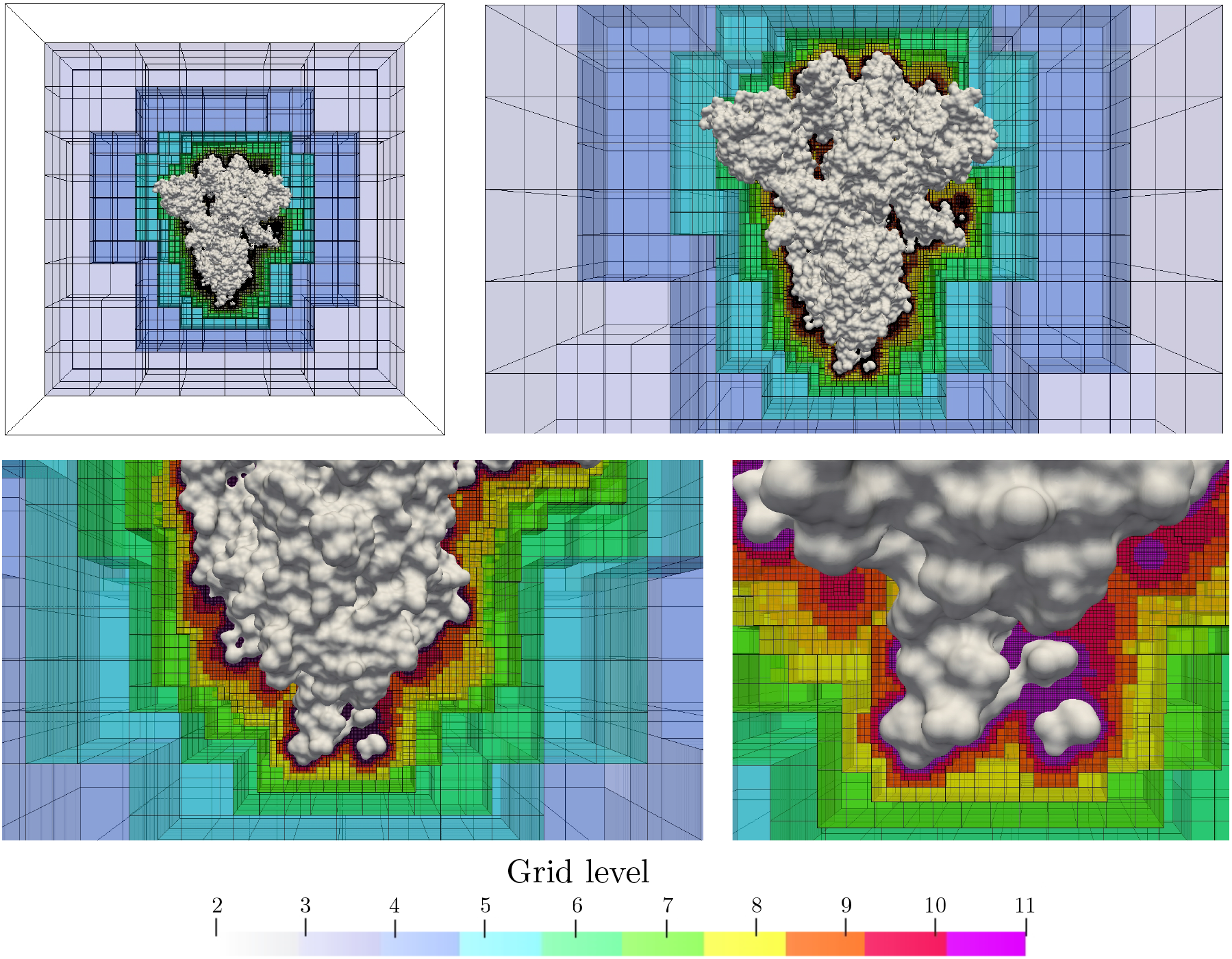
Vizualizations of the entire computational domain, SARS-CoV-2 spike protein (frame 1), and finest mesh used for this study (level 11 with 34, 362, 796 nodes). Computational cells are colored by their corresponding tree level (*i.e*. the number of successive mesh subdivisions required for their construction). For visual purposes only half of the mesh is depicted.

The entire computational domain is defined as the cube of side length 400Å (about twice the size of the whole protein) and center 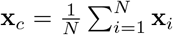. Calculations were performed on Octrees of maximum level ranging from 7 to 11. On the finest grids, the minimal spatial resolution is 0.18Å.

#### 4.2.2. Error Estimators

To monitor the convergence of the overall method we construct *a posteriori* numerical error estimators. They only rely on the decomposition presented in section 4.1.4, and can therefore be employed independently of the numerical approach. For the current implementation, detailed in section 4.2, we refer the interested reader to [26, 27] for formal convergence studies, using analytic solutions and order estimations.

The error on the interface representation, gradient of the solution (*i.e*. electric field) and solution itself can be estimated using the following metrics *e_Γ_,e_E_,e_ψ_*

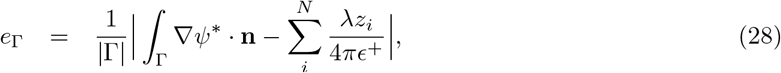

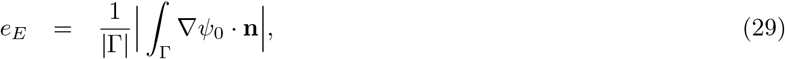

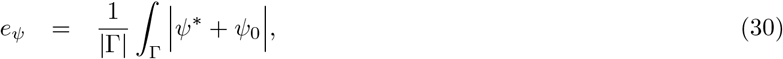

where |Γ| denotes the surface area of Γ. In virtue of Gauss’s Theorem the first two integrals are null. Because of the boundary condition (23), the third quantity should also be null. When computing *e*_Γ_, because the gradient of the Coulombic and the total charge are known exactly, numerical errors can only arise from calculating the local normal **n** or approximating the surface integral. Therefore, this metric focuses on the geometry and its manipulation only.

The numerical errors in the gradient of the component *ψ*_0_ are the main source of errors in the second metric *e_E_*. Thus, it can be interpreted as a lower bound estimate for the average total normal electric field on the interface. The metric *e_ψ_*, the average error in *ψ*_0_ on the interface, can similarly be used to estimate the numerical error in the total potential on the interface.

#### 4.2.3. Convergence Study

Figure 8*a* depicts the spike protein’s electrostatic potential at the initial frame (closed configuration) as the mesh is refined. Positive electrostatic potential is colored in blue, neutral (no charge) in white, and the negative electrostatic potential is shown in red. The coarsest simulations (max_level_ = 7, 8) are only able to reproduce the general structure of the protein but fail at creating an accurate electrostatic map. As the maximal resolution reaches the characteristic atomic radius (*r*_min_ = 1Å), the finest geometrical features are correctly reproduced, leading to appreciably more accurate results (max_level_ = 9, minimal resolution 0.79AÅ). Further increasing the spatial resolution refines these molecular structures and the small scale potential variations even more (max_level_ = 10,11).

**Figure 8:**
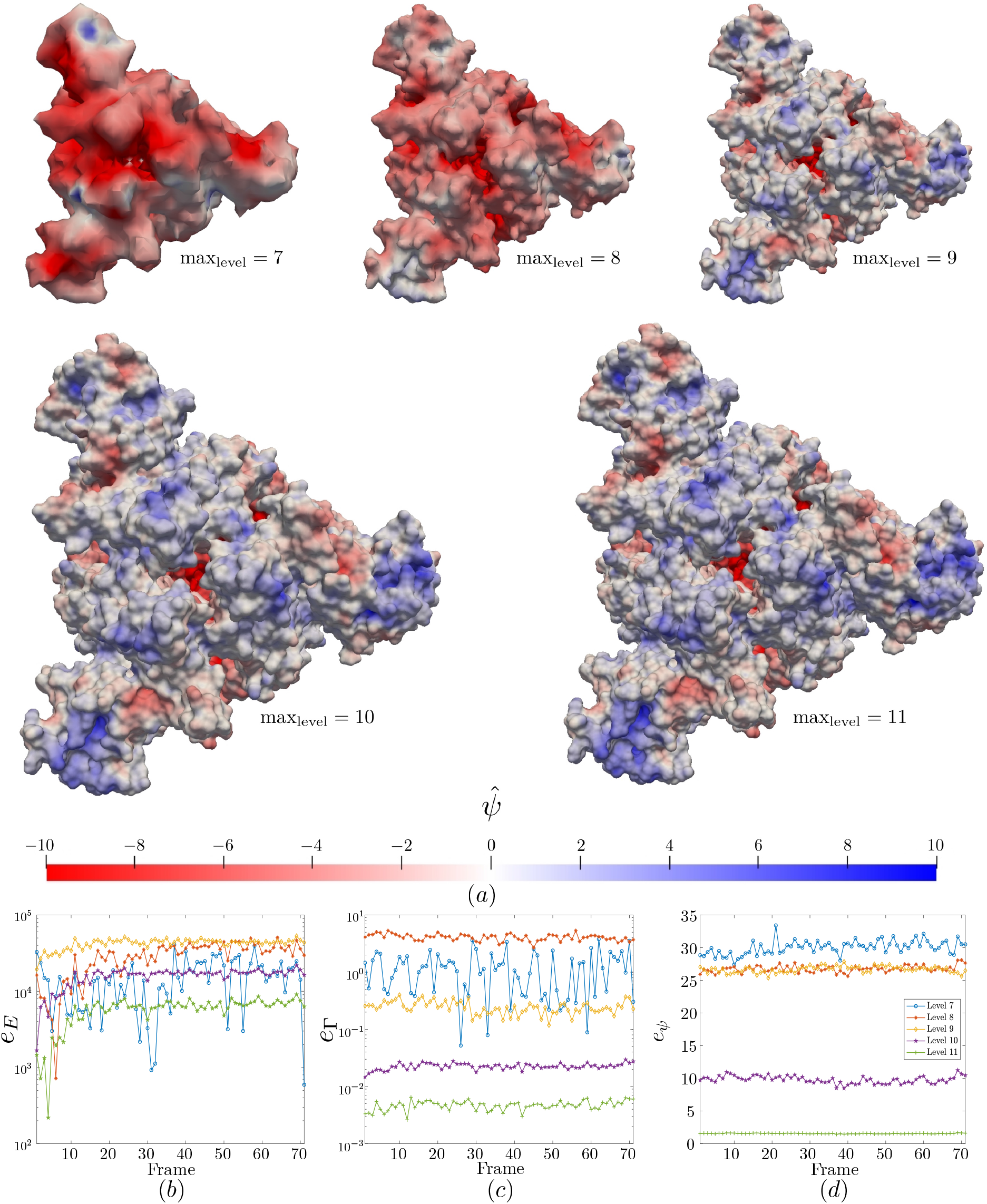
(*a*) Evolution of the spike’s Solvent Excluded Surface and electrostatic potential as the maximal resolution increases. (top down view in the closed configuration (frame 1)). (*b – d*) Convergence of the interface representation (*e*_Γ_), surface normal electric potential (*e_E_*), and surface potential (*e_ψ_*).

The time evolution of our three error estimates for all examined resolutions is presented in Fig. 8. As expected, all three metrics converge with increasing resolution. The impact of using subatomic resolution (*i.e*. max_level_ ≥ 9) is well illustrated with the convergence of the electric field and potential error estimates: it is unclear for super-atomic resolutions (max_level_ = 7, 8), and evident for subatomic ones (max_level_ = 9,10,11).

The error estimation for the total potential (*e_ψ_*) is significantly larger than the variations between consecutive maximum levels observed in Fig. 8a, which is a proxy for the error on the regular part of the solution 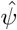. Since *e_ψ_* involves the singular part of the solution, which exhibits large spatial variations over small length scales, it is prone to higher numerical errors and is expected to be larger than the actual error on the regular part of the solution. Closer inspection of our measurements reveals that these two metrics may differ by at least one order of magnitude. The error estimation for *ψ*, using max_level_ = 10 is approximately equal to the maximum absolute surface potential value (≈ 10) observed on Fig. 8a. This misleadingly suggests the relative error on the total solution is as large as 100%. The comparison between the potential maps for max_level_ = 10 and max_level_ = 11 indicates that in practice the relative error on the surface potential probably lies between 1% and 10%.

From all these remarks, we are confident that the method is correctly implemented and that the most refined simulations accurately capture the S protein’s electrostatic potential. For the finest resolution, the estimated average error on the total potential is close to 2, which in light of the above discussion suggests that the average practical relative error on the surface potential is under 2%.

## 5. Acknowledgments

The authors would like to thank M. Zimmerman, G. Bowman, and the Folding@home project for creating and providing us with the S protein opening trajectory. We also thank D. Strubbe and J. Grasis for valuable discussions. This research was supported by a COVID-19 seed grant from the Center for Information Technology Research in the Interest of Society (CITRIS) at UC Merced awarded to M. Theillard and S. Sukenik. The authors acknowledge computing time on the Multi-Environment Computer for Exploration and Discovery (MERCED) cluster at the University of California, Merced, which was funded by National Science Foundation Grant No. ACI-1429783. M. Theillard and S. Sukenik are members of the NSF-CREST Center for Cellular and Biomolecular Machines at the University of California, Merced (NSF-HRD-1547848).

## 6. Author Contributions

M. Theillard and S. Sukenik conceived the presented project. M. Theillard developed and implemented the numerical method. S. Strango, A. Kucherova, and M. Theillard carried the computations and analyzed the Results. All authors contributed to the redaction of the manuscript and approved its final form.

